# The utility of native MS for understanding the mechanism of action of repurposed therapeutics in COVID-19: heparin as a disruptor of the SARS-CoV-2 interaction with its host cell receptor

**DOI:** 10.1101/2020.06.09.142794

**Authors:** Yang Yang, Yi Du, Igor A. Kaltashov

**Affiliations:** Department of Chemistry, University of Massachusetts-Amherst, 240 Thatcher Way, Amherst, MA 01003

## Abstract

The emergence and rapid proliferation of the novel coronavirus (SARS-CoV-2) resulted in a global pandemic, with over six million cases and nearly four hundred thousand deaths reported world-wide by the end of May 2020. A rush to find the cures prompted re-evaluation of a range of existing therapeutics vis-à-vis their potential role in treating COVID-19, placing a premium on analytical tools capable of supporting such efforts. Native mass spectrometry (MS) has long been a tool of choice in supporting the mechanistic studies of drug/therapeutic target interactions, but its applications remain limited in the cases that involve systems with a high level of structural heterogeneity. Both SARS-CoV-2 spike protein (S-protein), a critical element of the viral entry to the host cell, and ACE2, its docking site on the host cell surface, are extensively glycosylated, making them challenging targets for native MS. However, supplementing native MS with a gas-phase ion manipulation technique (limited charge reduction) allows meaningful information to be obtained on the non-covalent complexes formed by ACE2 and the receptor-binding domain (RBD) of the S-protein. Using this technique in combination with molecular modeling also allows the role of heparin in destabilizing the ACE2/RBD association to be studied, providing critical information for understanding the molecular mechanism of its interference with the virus docking to the host cell receptor. Both short (pentasaccharide) and relatively long (eicosasaccharide) heparin oligomers form 1:1 complexes with RBD, indicating the presence of a single binding site. This association alters the protein conformation (to maximize the contiguous patch of the positive charge on the RBD surface), resulting in a notable decrease of its ability to associate with ACE2. The destabilizing effect of heparin is more pronounced in the case of the longer chains due to the electrostatic repulsion between the low-p*I* ACE2 and the heparin segments not accommodated on the RBD surface. In addition to providing important mechanistic information on attenuation of the ACE2/RBD association by heparin, the study demonstrates the yet untapped potential of native MS coupled to gas-phase ion chemistry as a means of facilitating rational repurposing of the existing medicines for treating COVID-19.

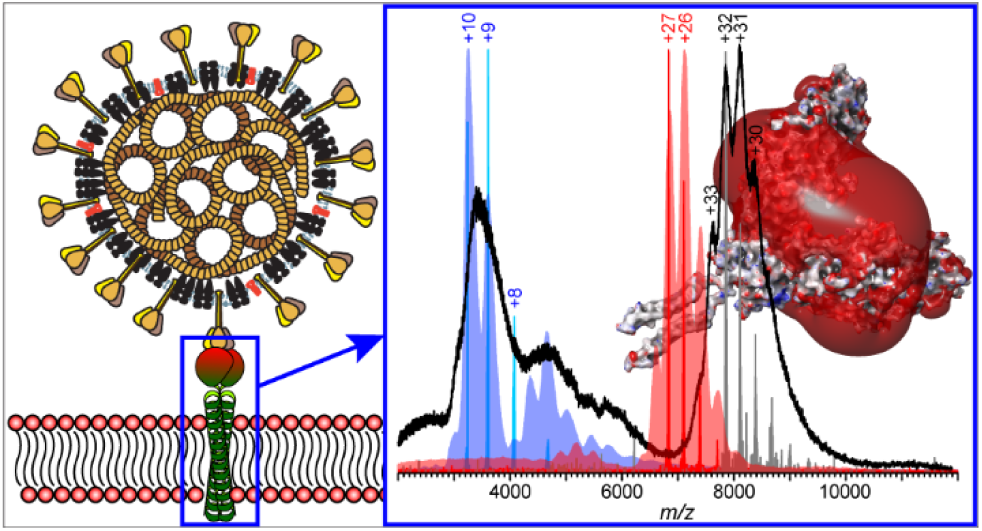

## INTRODUCTION

The emergence of the novel coronavirus (SARS-CoV-2) in late 2019^1^ resulted in a global pandemic that had left virtually no country in the world unaffected.^2^ The new disease (termed COVID-19) claimed over 400,000 lives worldwide by the end of May 2020, with the number of new cases still averaging over 100,000 in early June. This global healthcare crisis has resulted in a rush to find effective treatments for COVID-19, with strategies relying on repurposing of the existing medicines given high priority.^3^ While the initial efforts were largely empirical,^4,5^ the rapid progress in understanding the etiology of COVID-19 and accumulation of the vast body of knowledge on the SARS-CoV-2 life cycle and its mechanism of infectivity provided an extensive list of therapeutic targets for rational intervention.^6^ One such high-value target is the viral spike protein (S-protein),^7^ which is critical for both docking of the viral particle to its host cell surface receptor ACE2,^8^ and the concomitant fusion with the cell membrane followed by the delivery of the viral load.^9^

One particularly promising avenue for therapeutic intervention that currently enjoys considerable attention is blocking the ACE2/S-protein interaction site with either antibodies or small molecules.^10^ In particular, heparin interaction with the S-protein has been shown to induce conformational changes within the latter^11^ and to have inhibitory effects on the cellular entry by the virus.^12^ Combined with the well-documented anti-coagulant and anti-inflammatory^13^ properties of heparin (that are highly relevant vis-à-vis the two hallmarks of COVID-19, the coagulopathy^14,15^ and the cytokine storm^16^), this led to a suggestion that heparin or related compounds may play multiple roles in both arresting the SARS-CoV-2 infection and mitigating its consequences.^17,18^ In fact, heparin treatment of COVID-19 patients has been adopted by some physicians and is associated with a better prognosis.^19^ At the same time, the use of heparin raises the specter of heparin-induced thrombocytopenia (HIT), and its incidents were found to be particularly high among critical COVID-19 patients.^20^ Clearly, utilization of heparin or related compounds as a safe and efficient treatment of coronavirus-related pathologies will hinge upon the ability to select a subset of structures that exhibit the desired properties (e.g., the ability to block the ACE2/S-protein association) while lacking the deleterious effects (e.g., the ability to create immunogenic ultra-large complexes with platelet factor 4, the hallmark of HIT,^21^ or cause excessive bleeding). Similar sentiments can be expressed with respect to a wide range of other medicines that are currently a focus of extensive repurposing efforts.^3^ This work can be greatly facilitated by analytical methods capable of providing detailed information on the drug candidates’ interactions with their therapeutic targets, and their ability to disrupt the molecular processes that are critical for the SARS-CoV-2 lifecycle. Native mass spectrometry (MS) has been steadily gaining popularity in the field of drug discovery,^22,23^ but its applications are frequently limited to relatively homogeneous systems. Unfortunately, the large size and the extensive glycosylation of the proteins involved in the SARS-CoV-2 docking to the host cell surface (fourteen N-glycans within the ectodomain of ACE2 and at least eighteen O- and N-glycans within the S-protein ectodomain,^24^ including three in its receptor binding domain, RBD) make the straightforward application of native MS to study ACE2/S-protein association challenging. This problem may be further exacerbated by the structural heterogeneity of therapeutics that are evaluated as potential disruptors of the ACE2/S-protein association, such as heparin and its derivatives.

Several approaches have been developed in the past decade as a means of facilitating native MS analyses of highly heterogeneous systems, which rely on supplementing MS measurements with non-denaturing front-end separation techniques,^25^ and gas-phase chemistry (e.g., limited charge reduction^26^). The latter is particularly attractive, as it allows native MS to be applied to systems as heterogeneous as associations of proteins with unfractionated heparin.^27^ In this work we use native MS in tandem with limited charge reduction to characterize ACE2/RBD complexes and evaluate the influence of heparin-related compounds (a synthetic pentasaccharide fondaparinux and a fixed-length eicosasaccharide heparin chains) on the stability of these complexes. Above and beyond providing important mechanistic details on attenuation of the ACE2/RBD association by heparin, the study demonstrates the potential of native MS supplemented by limited charge reduction to support the COVID-19 related drug repurposing efforts.

## EXPERIMENTAL SECTION

The recombinant forms of human ACE2 (residues 1-740) and RBD (residues 319-541) were purchased from Sino Biological (Wayne, PA). Both proteins were extensively dialyzed in 150 mM NH_4_CH_3_CO_2_ prior to MS analyses. Fondaparinux was purchased from Sigma-Aldrich (St. Louis, MO), and heparin eicosasaccharide produced by partial depolymerization of heparin was purchased from Iduron (Alderley Edge, UK). All solvents and buffers used in this work were of analytical grade or higher.

Native MS measurements were carried out using a Synapt G2-Si (Waters, Milford, MA) hybrid quadrupole/time-of-flight mass spectrometer equipped with a nanospray ion source. The following ion source parameters were used to maintain non-covalent complexes in the gas phase: sampling cone voltage, 80 V; trap CE, 4 V; trap DC bias, 3 V; and transfer CE, 0 V. Ion selection before limited charge reduction was carried out by setting the quadrupole selection parameters (LM resolution) to 4.5. To trigger limited charge reduction, the trap wave height was set at 0.3 V and the discharge current was optimized.

Molecular modeling of the RBD/heparin complexes was carried out using a Maestro (Schrödinger LLC, New York, NY) modeling suite, release 2019-4. The model of RBD/ACE2 was prepared using the 2019-nCoV RBD/ACE2-B0AT1 complex (PDB 6M17) as a template. The pentasaccharide model was extracted from the crystal structure of the platelet factor 4/fondaparinux complex (PDB 4R9W) and dp20 model was created by deleting a tetrasaccharide from the nonreducing end of heparin dp24 (PDB 3IRJ) with 3 sulfate groups cut-off. The heparinoid complexes were minimized using the OPLS3 force field. The MD simulations were set up using a neutralized system (with 6 and 34 Na+ ions used for fonfdaparinux and dp20, respectively) in explicit water and 150 mM NaCl at 300K under the OPLS3 force field.

## RESULTS AND DISCUSSION

Despite the modest size of the SARS-CoV-2 receptor binding domain (RBD), its mass spectrum acquired under native conditions (Figure 1, bottom) is convoluted and difficult to interpret. Since the main source of structural heterogeneity within this protein is its extensive glycosylation (Shajahan, et al. identified two N-glycosylation and at least one O-glycosylation sites within this segment of the S-protein^24^), we also attempted to work with a commercial RBD expressed in *E. coli*, which presumably lacked glycan chains. However, this construct had poor solubility characteristics and appeared to exist as a large aggregate (mostly likely due to the aberrant disulfide formation). Therefore, our efforts were focused on using limited charge reduction to facilitate the native MS analysis of glycoproteins expressed in eukaryotic cells, which identified the major ionic species as the RBD monomer (average MW 32.7 kDa), and the minor one as the dimer (65.5 kDa). The latter is likely to reflect the presence of an unpaired cysteine residue within the RBD segment (see *Supporting Information*) capable of forming an external disulfide bond.

**Figure 1.**
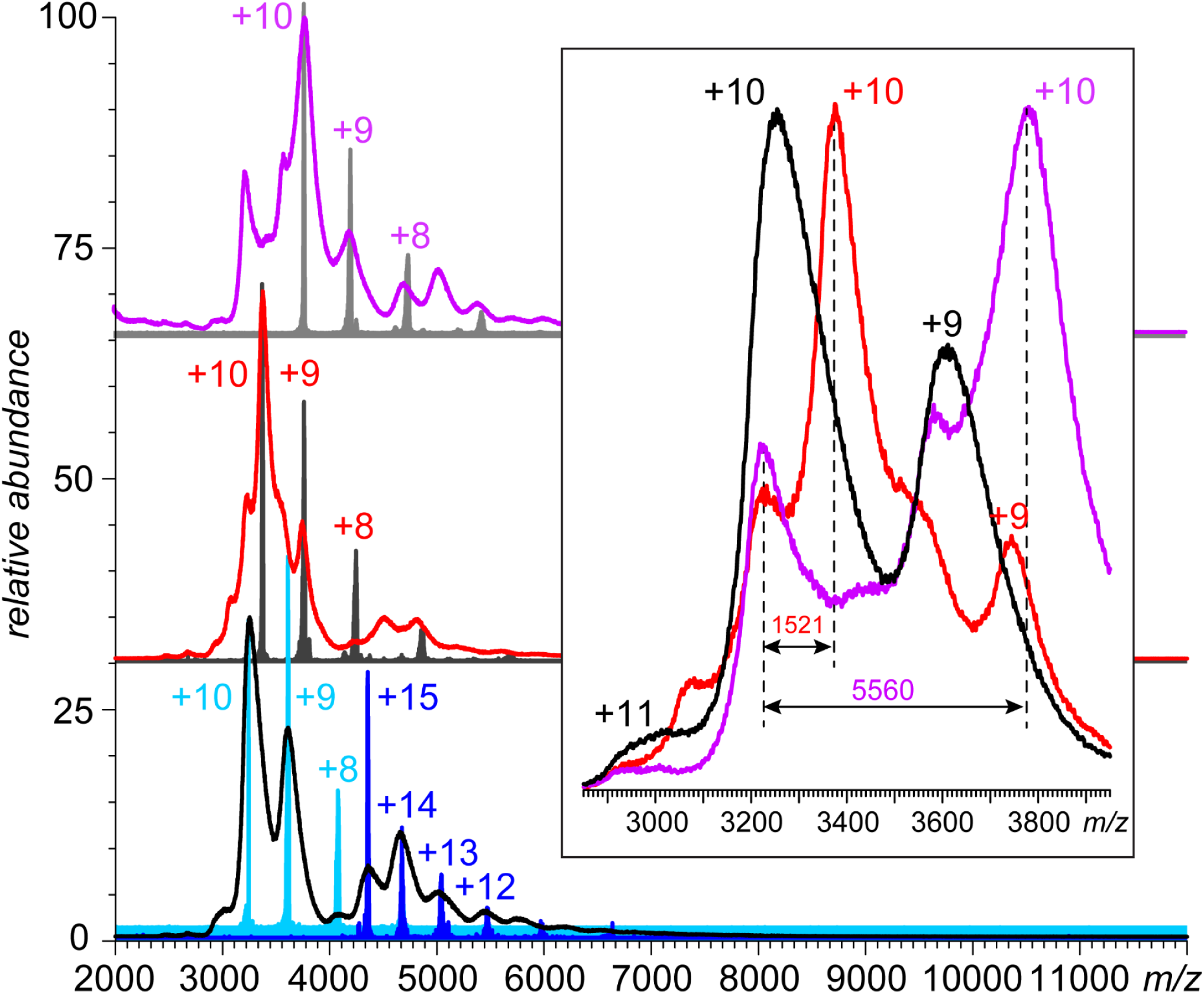
Mass spectra of the recombinant form of the SARS-CoV-2 S-protein RBD (10 μM aqueous solution in 150 mM NH_4_CH_3_CO_2_) in the absence of heparinoids (bottom) and in the presence of 15 μM fondaparinux (middle) and 15 μM fixed-length heparin oligomer dp20 (top). The color-filled spectra in each panel show the charge ladders generated by the limited charge reduction measurements. The inset shows a zoomed view of the ionic signals at the charge state +10 from all three mass spectra.

Addition of a structurally homogeneous pentasaccharide heparin mimetic (fondaparinux) to the RBD solution results in a noticeable shift of the ionic signal in native MS (Figure 1, middle). The magnitude of this shift (1521 Da) correlates with the ligand mass (1505 Da); importantly, only 1:1 protein/ligand complexes are observed (in contrast to high-pI proteins, which act as heparin “sponges” by accommodating multiple polyanionic chains^28^). Similar behavior was observed when a fixed-length heparin oligomer (eicosasaccharide, or dp20) was added to the RBD solution. The magnitude of the shift (5.6 kDa) corresponded to a dp20 species carrying on average 26 sulfate groups (the sulfation levels in dp20 range from 17 to 28^29^), and only 1:1 RBD/dp20 complexes were observed alongside the less abundant free protein (see the inset in Figure 1).

The absence of the 1:2 protein/heparin oligomer complexes might seem surprising, as the protein surface contains two distinct basic patches (R346, R355, K356 and R357) and (R454, R457, K458, K462 and R466) that were previously suggested to be heparin binding sites.^11^ Molecular dynamics (MD) simulations of the RBD/fondaparinux complex indicates that the polyanion association with the protein results in significant conformational changes on the surface of the latter, giving rise to a consolidated patch of the positive charge (Figure 2). While such conformational rearrangement provides enthalpic gains for the electrostatically driven RBD/heparin oligomer interaction, it exerts deleterious effects on the RBD/ACE2 binding. Indeed, conformation of the receptor-binding motif (RBM) of RBD following RBD association with fondaparinux undergoes significant changes, which affect critical residues in the RBD/ACE2 interface^7^ (see *Supporting Information* for more detail).

**Figure 2.**
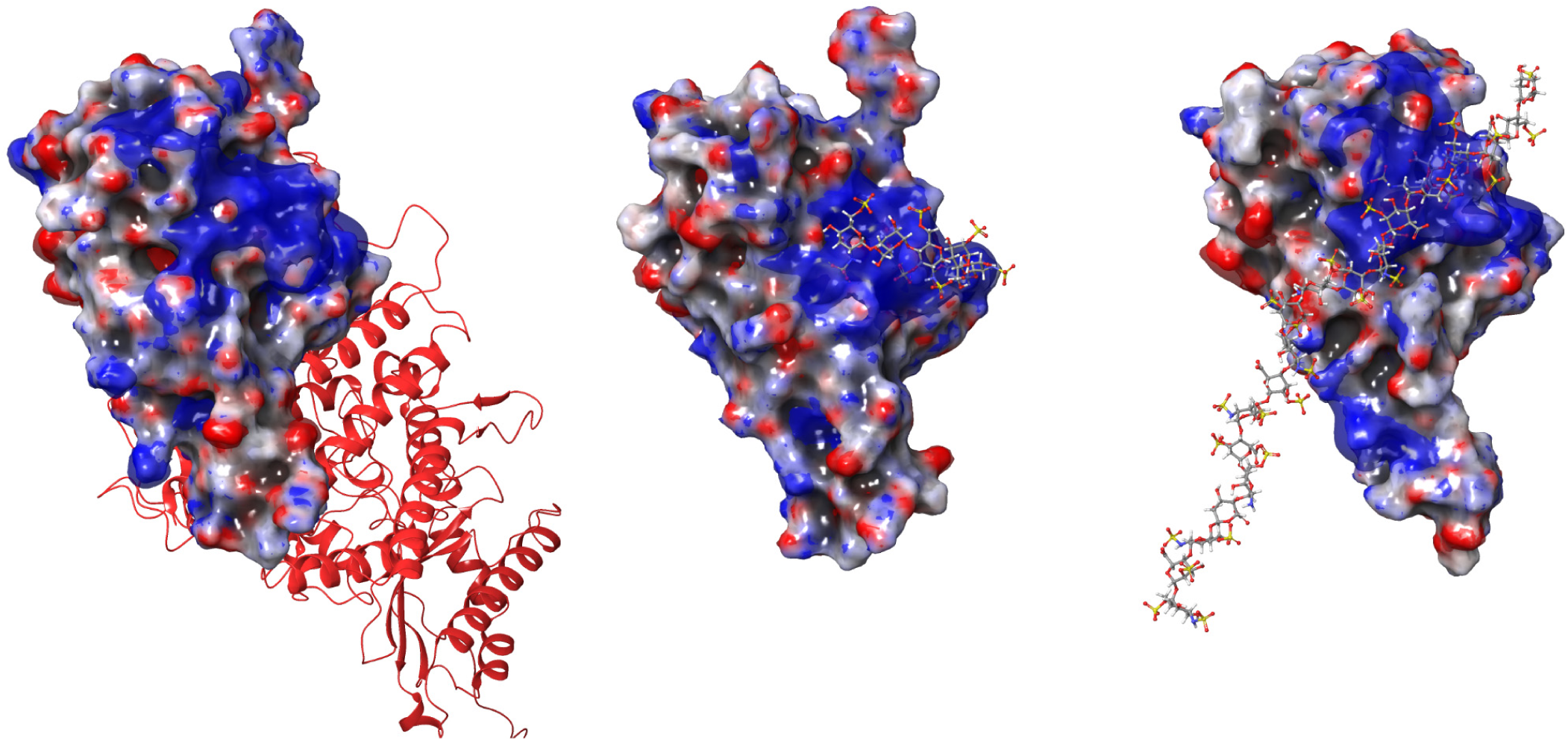
The 3*kT*/*e* isopotential surfaces calculated for RBD associated with ACE2 (left), fondaparinux (middle) and dp20 (right). The RBD/ACE2 structure (part of pbd 6M17^31^) shows ACE2 in a ribbon format (red). Both RBD/heparinoid complexes show representative structures obtained from MD simulations; both heparinoids are shown in a ball-and-stick format (oxygen and sulfur atoms are colored in red and orange, respectively).

To confirm the ability of a short heparin oligomer to disrupt the RBD/ACE2 association, native mass spectra of the two proteins’ mixture were acquired in the absence and in the presence of fondaparinux. The reference RBD/ACE2 spectrum (Figure 3, bottom) features an abundant signal of the (RBD°ACE2)_2_ complex alongside the residual (unbound) RBD that was present in molar excess in solution. No signal of unbound ACE2 could be detected, consistent with the reported binding strength in the low-nM range.^7^ The mass spectrum appearance changes in the presence of fondaparinux (5-fold molar excess over RBD), with the ionic signal of ACE2 monomers becoming prominent in the *m/z* region 5,000-6,000 (Figure 3, middle trace). The presence of both free RBD and ACE2 species suggests a dramatic decrease of the affinity (to the μM level). The destabilizing effect is even more significant in the presence of the longer heparinoid. The presence of dp20 in solution resulted in the ionic signals of free RBD and ACE2 monomer becoming equiabundant with that of their 2:2 complex; the spectrum also reveals the presence of an unsaturated (2:1) ACE2/RBD complex (Figure 3, top).

**Figure 3.**
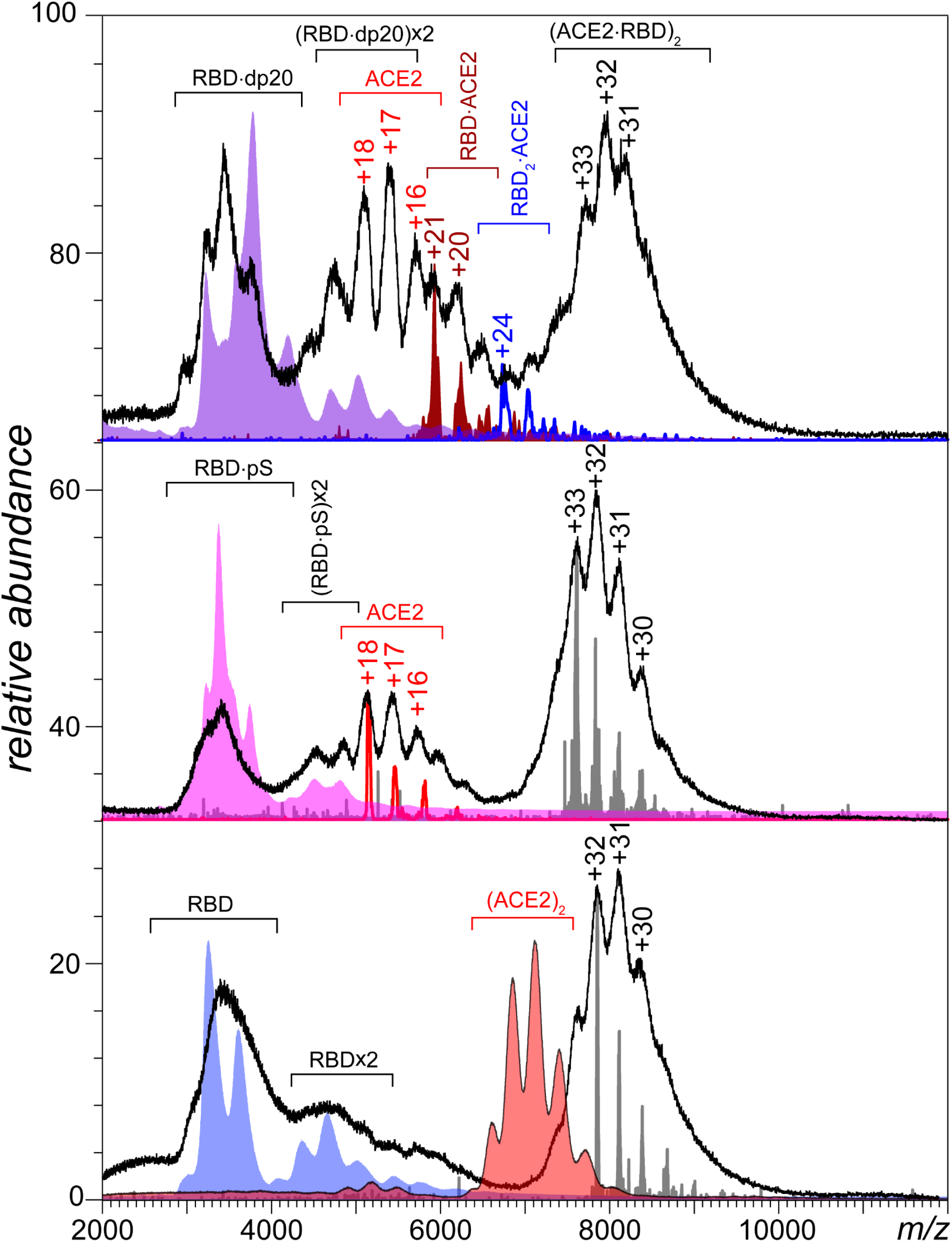
Native MS of RBD/ACE2 solutions (5 and 2.5 μM, respectively, in 150 mM NH_4_CH_3_CO_2_) acquired in the absence of heparinoids (bottom), and in the presence of fondaparinux (middle) and dp20 (top). The faded color-filled curves show reference mass spectra of ACE2 (red), RBD (blue), RBD/fondaparinux (pink) and RBD/dp20 (purple). The well-defined charge ladders show the results if limited charge reduction measurements that were used to assign charges to poorly-defined ion peaks in the native MS.

The importance of the heparin oligomer chain length in modulating its ability to disrupt the RBD/ACE2 interaction is likely to be related to the limited physical size of the positive patch on the RBD surface that cannot accommodate more than six saccharide units (Figure 2). The rest of the polyanionic chain is exposed to the solvent, resulting in unfavorable interactions with the low-pI ACE2 molecules (due to the electrostatic repulsion). This results in a synergistic effect, with the longer heparin chain destabilizing the RBD/ACE2 association via both conformational rearrangements within the former (*vide supra*) and the long-range electrostatic repulsion of the latter. In contrast, the shorter oligomer (fondaparinux) disrupts the RBD/ACE2 interaction only via allosteric conformational changes within the RBM segment, exerting a somewhat weaker destabilizing effect. Taken together, the results of native MS measurements and MD simulations provide important insights into the mechanism of the RBD/ACE2 association disruption by heparin that will be invaluable for rational selection of the most potent inhibitors of the SARS-CoV-2 docking to the host cell.

## CONCLUSIONS

The level of structural heterogeneity displayed by both viral proteins and their counterparts on the surface of the host cells (as well as some of the proposed therapeutics, such as heparin) may seem overwhelming for the straightforward “intact-molecule” MS measurements. However, incorporation of the limited charge reduction in the experimental workflow allows meaningful information to be obtained on objects as complex as 2:2 RBD/ACE2 associations. While native MS cannot provide the level of structural detail produced in crystallographic studies,^30,31^ it allows the influence of various compounds on the stability of such complexes to be readily evaluated and mechanistic details to be revealed, thereby enabling rational approach to the drug repurposing efforts. Design of successful therapeutic strategies against a foe as formidable as SARS-CoV-2 will require mobilization of efforts and resources in the entire field of life sciences, and the analytical tools (including MS) will undoubtedly play a pivotal role in these efforts.

## ASSOCIATED CONTENT

### Supporting Information

Supporting Information includes (*i*) ion source parameters used in MS measurements; (*ii*) ACE2 and RBD amino acid sequences; (*iii*) native MS of ACE2; and (*iv*) details of MD studies of the heparin oligomers interactions with RBD.

## Author Contributions

Y.Y. designed and carried out the experimental work, and analyzed the data. Y.D. carried the out molecular modeling work. I.K. designed the study, analyzed the data and drafted the manuscript. All authors have given approval to the final version of the manuscript.

## ACKNOWLEDGMENT

This work was supported by grants from the National Institutes of Health (R01 GM112666) and the National Science Foundation (CHE-1709552). MS instrumentation used in this work is part of the Mass Spectrometry Core facility at UMass-Amherst.

## Supplementary Information for

### Ion source parameters used in MS measurements

The following ion source parameters were used to maintain non-covalent complexes in the gas phase: sampling cone voltage, 80 V; trap CE, 4 V; trap DC bias, 3 V; and transfer CE, 0 V. Ion selection before limited charge reduction was carried out by setting quadrupole selection parameters: LM resolution, 4.5. To trigger limited charge reduction, the trap wave height was set as 0.3 V and the discharge current was optimized.

### ACE2 amino acid sequence

The sequence is based on the UniProt entry Q9BYF1. The black vertical arrows indicate the termini of the construct used in this work (not inclusive of the C-terminal His-tag). The SARS-CoV-2 S-protein RBD contact sites are highlighted in blue. The presumed N-glycosylation sites are underlined, and the disulfide bonds are indicated with brackets. The gray-shaded region (not included in the construct) is the transmembrane part of ACE2.

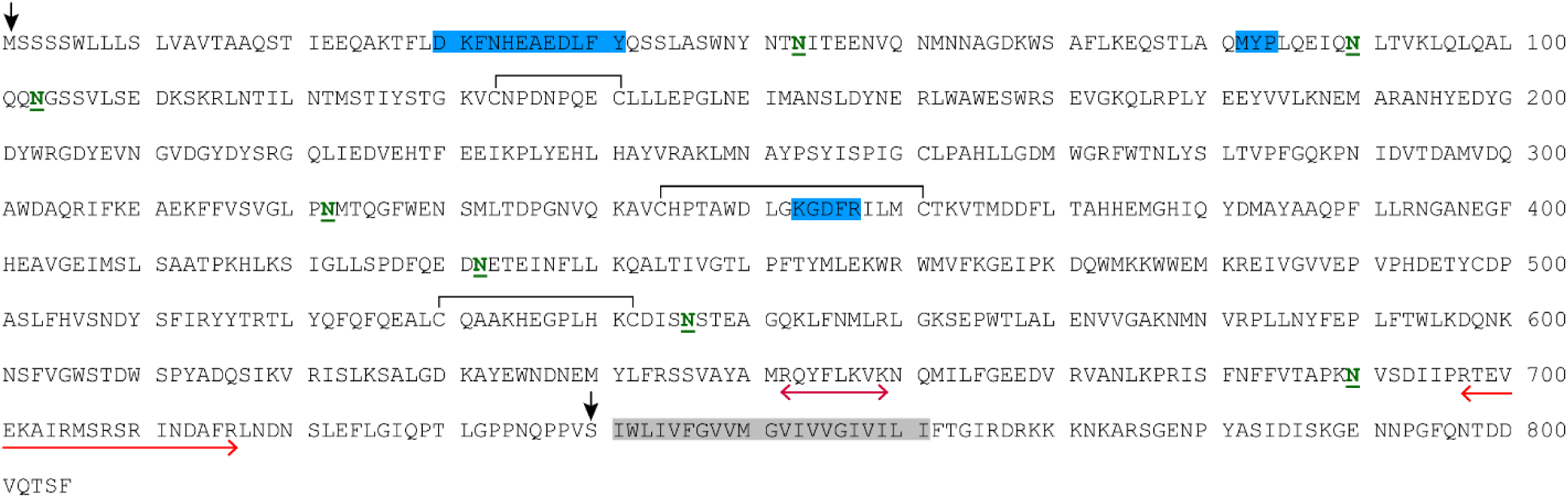

### RBD amino acid sequence

The RBD sequence (underlined with blue lines) is shown in the context of UniProt entry P0DTC2 representing the entire ectodomain of the SARS-CoV-2 S-protein (the gray-shaded region corresponds to the S1 domain). The amino acid residues forming distinct positive patches on the RBD surface are colored in blue. The presumed N-glycosylation sites are underlined, and the N- and O-glycosylation sites reported by Shajahan, et al. (<u>Deducing the N- and O-glycosylation profile of the spike protein of novel coronavirus SARS-CoV-2</u>. *Glycobiology* **2020**, in press) are indicated with the blue and yellow boxes, respectively (the blue crosses indicate the presumed glycosylation sites that were reported to be glycan-free by Shajahan, et al.). The disulfide bonds are indicated with black brackets. The red-shaded regions indicate the protease cleavage sites. The fusion peptide and the heptad repeats are underlined with green and brown lines, respectively.

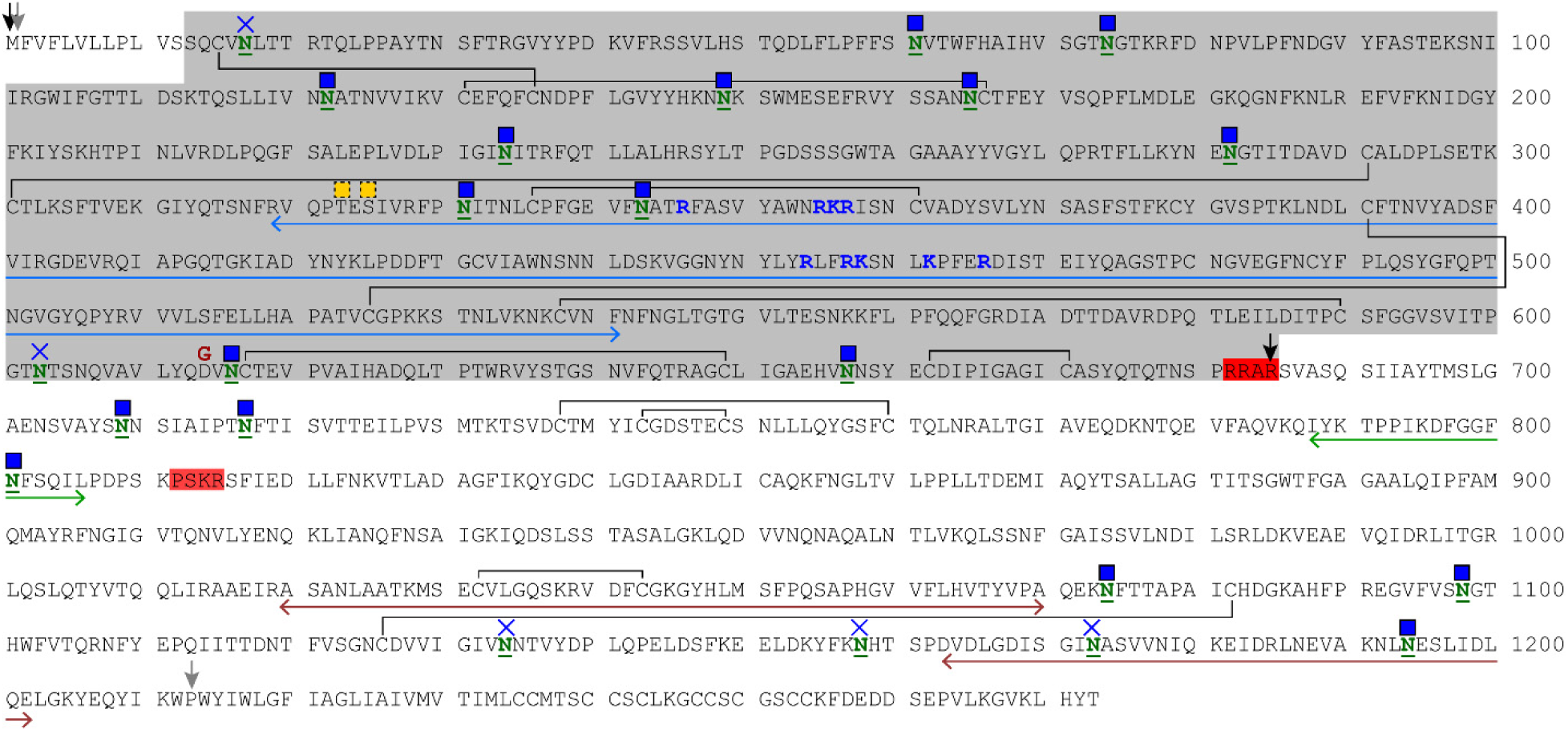

### Native MS of ACE2

Black trace: a mass spectrum of an aqueous solution of ACE2 ectodomain (5 μM in 150 mM NH_4_CH_3_CO_2_, pH 6.9). Green, orange and red traces: signal generated by limited charge reduction of ionic populations selected at *m/z* 5,161 u, 6,600 u and 6,855 u, respectively (the masses deduced from these charge ladders are 92,954 Da and 185,264 Da, representing the monomeric and dimeric states of ACE2 ecto-domain)

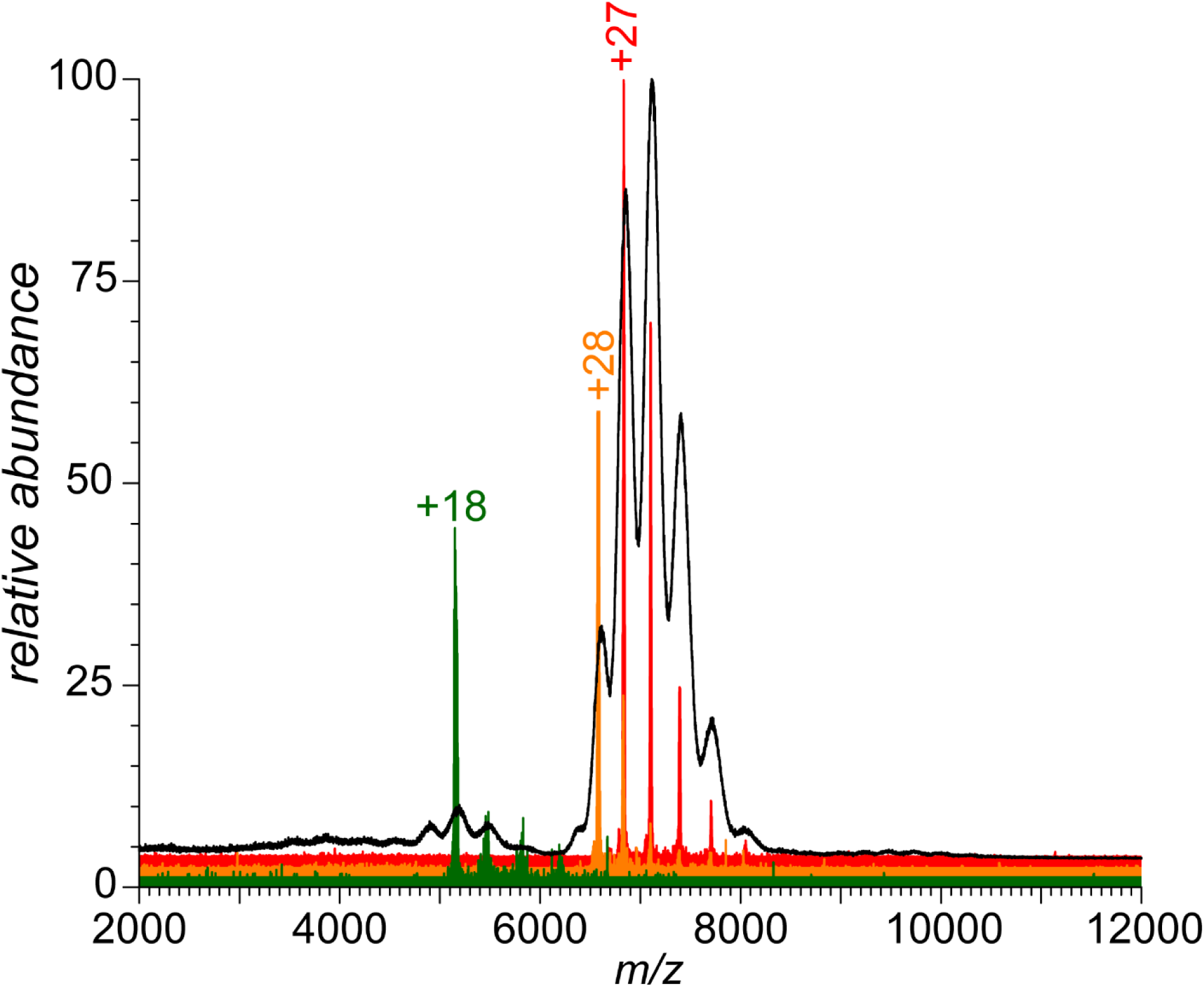

### Details of MD studies of the heparin oligomers interactions with RBD

The model of RBD/ACE2 was prepared using the 2019-nCoV RBD/ACE2-B0AT1 complex (PDB 6M17) as a template. The pentasaccharide model was extracted from platelet factor 4/fondaparinux (PDB 4R9W) and dp20 model was created by deleting a tetrasaccharide from the nonreducing end of heparin dp24 (3IRJ) with 3 sulfate groups cut-off. The heparinoid complexes were minimized using the OPLS3 force field. The MD simulations were set up using a neutralized system (with 6 and 34 Na+ ions used for fonfdaparinux and dp20, respectively) in explicit water and 150 mM NaCl at 300K under OPLS3 force field.

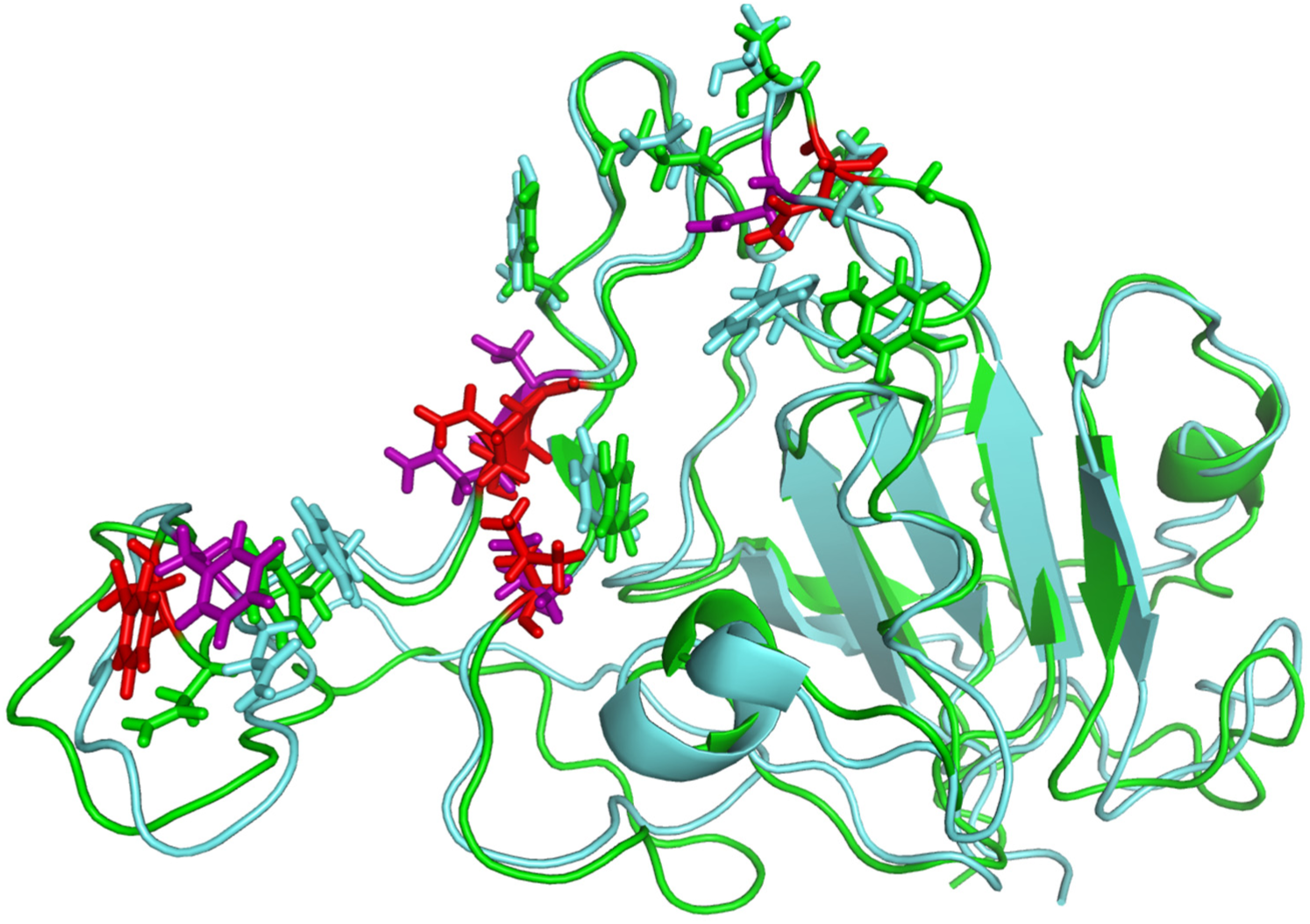

Influence of fondaparinux binding to RBD on the conformation of the ACE2-binding interface. The ribbon diagrams show backbones of RBD bound to ACE2 (green) and associated with fondaparinux following 1.5 ns MD simulation (teal). Side chains of all contact residues are shown in stick representation; the five critical residues are colored in each structure (red and purple, respectively).

